# Correlating membrane-protein dynamics with function: Integrating bioinformatics, molecular dynamics, and single-molecule FRET

**DOI:** 10.1101/2025.06.05.658054

**Authors:** Hugh R. Higinbotham, Christine A. Arbour, Barbara Imperiali

**Author notes:** **Corresponding author**: Barbara Imperiali - Department of Biology and Department of Chemistry, Massachusetts Institute of Technology, Cambridge, MA 02139. Hugh R. Higinbotham - Department of Biology and Department of Physics, Massachusetts Institute of Technology, Cambridge, MA 02139;, Christine A. Arbour – Department of Biology and Department of Chemistry, Massachusetts Institute of Technology, Cambridge, MA 02139; Present Address: Synthetic Molecule Design and Development (SMDD), Eli Lilly and Company, Indianapolis, IN 46221. Detailed procedures for bioinformatic and MD analysis; protein expression and purification, fluorophore labeling, activity assays; slide functionalization and microscope setup; and smFRET data collection and image analysis. Supplementary videos: V1-6 of select PGTs ± ligands over 200 ns, aligned with membrane hidden; V7 of full 1 μs extended simulation of V2.

## Abstract

We present a strategy that deploys structural bioinformatics, molecular simulation, and single-molecule FRET microscopy for observing the ligand-dependent conformational dynamics of integral membrane proteins *in situ.* We focus on representative members of the small monotopic phosphoglycosyl transferase (SmPGT) superfamily, which catalyze transfer of a phosphosugar from a soluble nucleotide-sugar donor to a membrane-embedded polyprenol phosphate acceptor in the initiating step of glycoconjugate biosynthesis in prokaryotes. Substrate-specific structural features were identified across the superfamily and correlated with ligand-dependent conformational dynamics in all-atom simulations. To experimentally validate the role of this motion in ligand binding, we developed a platform to monitor intra-molecular protein dynamics in a native-like lipid environment. The presented approach incorporates selective cysteine protein labeling and non-canonical amino acid mutagenesis with bicyclononyne-tetrazine click chemistry to assemble dual-labeled variants of PglC, the initiating enzyme of the N-linked protein glycosylation pathway from *Campylobacter jejuni.* The modified proteins are then solubilized into styrene maleic acid liponanoparticles (SMALPs) to maintain an *in situ* membrane environment. The conformational changes of PglC upon inhibitor binding are diagnostic of inhibitor potency. The single-molecule FRET-SMALP strategy can be adapted to investigate protein dynamics across the superfamily of SmPGTs with different substrate selectivity where structure prediction and molecular dynamics support significant conformational changes upon ligand binding.

**Broader Impact:** Membrane protein structure-function relationships are critical for understanding fundamental biological processes and for the development of small-molecule drug treatments. Bacterial glycoconjugate biosynthesis pathways are a promising target for strain-specific antibiotics and exemplify the challenges of characterizing biomolecular systems that depend on highly specific protein, lipid, and carbohydrate chemistries. We integrate molecular simulation, structural bioinformatics, and single-molecule FRET to elucidate details of small-molecule binding to the PGT superfamily.

## Introduction

Integral membrane proteins (IMPs) comprise approximately 25% of proteomes across kingdoms of life (Krogh et al. 2001). The specific composition and physicochemical properties of membranes vary widely, and the diverse interplay of this environment with IMP structure and function is a rich target for investigation (Teo et al. 2019; Levental and Lyman 2023; Brown et al. 2025). Major advances in X-ray crystallography and Cryo-electron microscopy have led to the determination of many membrane protein structures (Nastou et al. 2020), and new membrane mimetic solubilization techniques are being introduced with increasing frequency (Autzen et al. 2019; Choy et al. 2021; Kermani 2021; Biou 2023; Ayub et al. 2024). Through the application of conformational restraints or advanced data processing, these techniques can also capture conformational variation in proteins in different “snapshots” and provide crucial information about their structural dynamics (Wu and Rapoport 2021; Zhong et al. 2021; Meszaros and Westenhoff 2024). However, despite these advances, the perturbations to IMPs necessary for sample preparation pose a challenge to connecting high-resolution structural information to the native dynamics associated with functional enzyme activity. Methods that simulate and experimentally characterize protein conformational dynamics in biologically relevant membrane environments are therefore crucial to link information from high-resolution structural studies to the native physiological context of IMPs.

IMPs in membrane-associated glycan assembly pathways are of particular interest due to their essentiality in bacterial survival, host-microbe interactions, and pathogenicity (Costa and Iraola 2019; Szymanski 2022). These pathways function at the cytoplasmic face of cell membranes and represent key targets for developing new classes of antibiotics that inhibit the biosynthesis of selected virulence-associated bacterial glycoconjugates. Critical to many glycoconjugate assembly pathways are phosphoglycosyl transferase (PGT) enzymes, which initiate glycan biosynthesis with a chemically distinct priming step. PGTs are categorized into two superfamilies distinguished by the membrane topology (monotopic or polytopic) of the minimal catalytic domain (O’Toole et al. 2021). The monotopic PGTs (monoPGTs) are exclusively prokaryotic and coordinate and catalyze phosphosugar transfer from a soluble nucleotide-sugar donor to a polyprenol phosphate acceptor that anchors the forming glycoconjugate to the membrane (Das et al. 2017a). Proteome-wide bioinformatics and functional analyses have recently enabled the classification of small monoPGTs, representing the minimal catalytic subunit, into clusters that share a common UDP-sugar substrate (Durand et al. 2024).

The best characterized small bacterial PGT is PglC from *Campylobacter concisus*; PglC initiates the general N-linked glycosylation pathway (Nothaft et al. 2012; Cain et al. 2020). The *Cc* PglC is the only small PGT (SmPGT) with a solved experimental structure. Variations between monomers in an X-ray structure that captures eight monomers in the octameric asymmetric unit and analysis of the position of mobile loop (residues 61-81) correlate with molecular dynamics (MD) simulation in all-atom bilayer systems. However, the connection between these structural dynamics and PGT function remain unclear (Ray et al. 2018; Majumder et al. 2023). The *Cc* PglC structure was recently updated to resolve additional C-terminal residues, and identifies a conserved aromatic box motif that anchors the C-terminus in place (Figure 1A) (Anderson et al. 2023). With both a validated full-length structure of *Cc* PglC and vastly expanded structural information across the superfamily of SmPGTs, we can now investigate structural dynamics across different UDP-sugar substrate specific families (Durand et al. 2024). Further investigation of the PGT conformational dynamics associated with substrate and ligand binding, conducted at the membrane interface, requires a targeted approach that integrates computational and experimental approaches.

**Figure 1:**
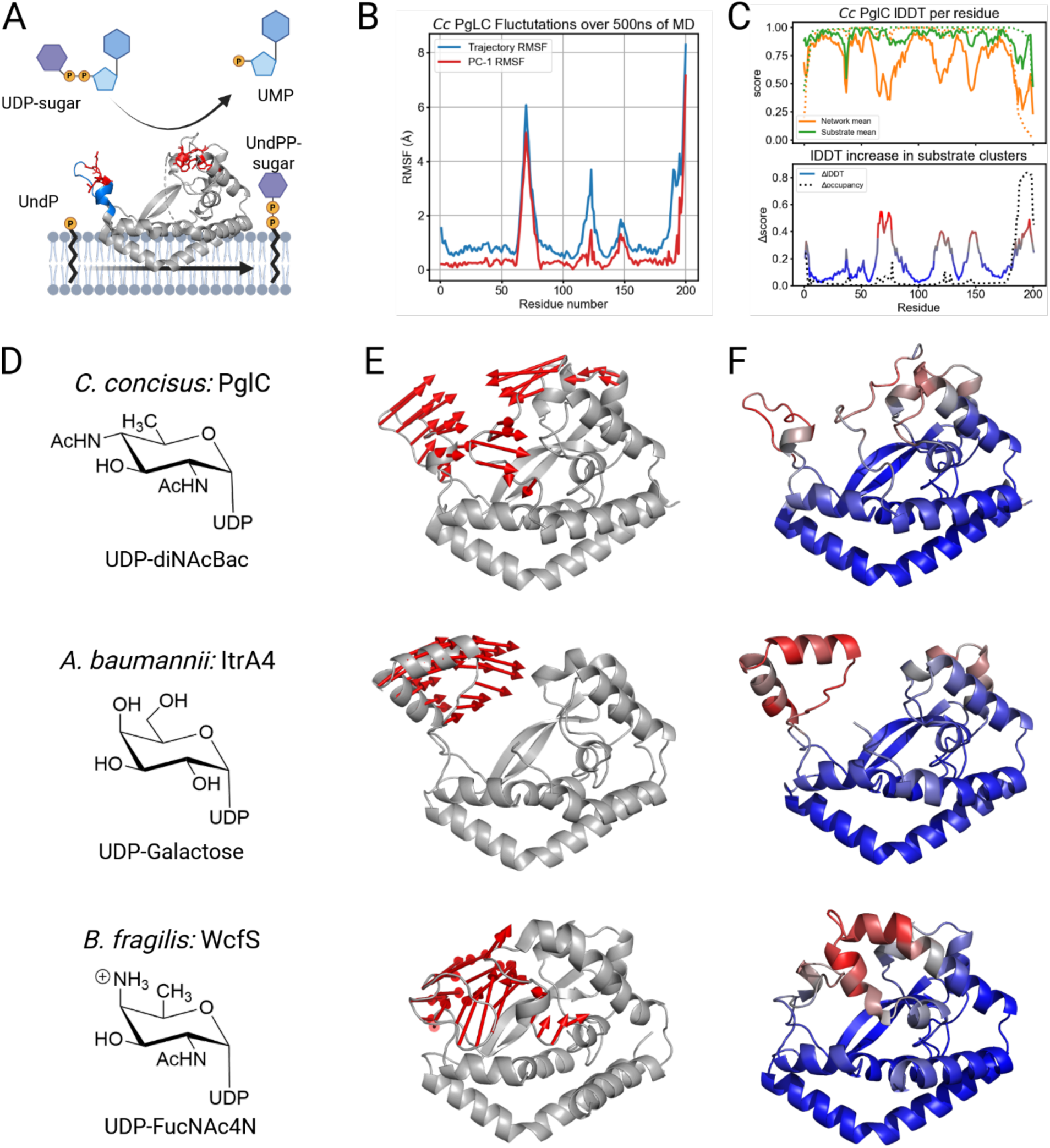
Structural dynamics of select SmPGTs. (A) Schematic of the PGT reaction with the crystal structure of Cc PglC (PDB: 8G1N). The substrate specific GLLP and aromatic box sequence motifs are highlighted in red, and blue indicates the mobile loop (residues 61-81). (B) RMSF by residue of Cc PglC over 500 ns of MD simulation. Red shows specifically the first principal component. (C) Top: average lDDT in pairwise structure alignments between Cc PglC and other PGTs in the SSN (Durand et al. 2024). Orange shows the network average and green shows only the diNAcBac cluster. Dotted lines show PGT occupancy at a given residue. Bottom: lDDT increase within substrate clusters by residue. Dotted line shows increase in occupancy. (D) PGT identity and cognate sugar substrate. (E) First principal component of PGT backbone over a 500 ns molecular dynamics trajectory in a membrane bilayer. (F) AlphaFold2 structure predictions colored by average lDDT increase in substrate specific clusters per residue.

Herein we report a combined computational and experimental strategy for investigating membrane protein dynamics in the presence of small molecule ligands to trigger and observe conformational changes (Zhang et al. 2015). We apply this approach to PglC from *Campylobacter jejuni* (*Cj*) and observe correlated closure of its mobile loop in the presence of nucleoside analog inhibitors that show improved binding. By combining information from structural bioinformatics, molecular dynamics simulations, and single-molecule Förster Resonance Energy Transfer (smFRET) microscopy, we assemble a clearer picture of PGT-substrate interactions, which will corroborate and aid the design of future structural studies and small molecule inhibitor discovery campaigns.

## Results

### Computational evidence of ligand-dependent loop mobility

We first sought to identify key dynamic features across the SmPGT superfamily. A given SmPGT catalyzes the transfer of a phospho-sugar to undecaprenol phosphate (UndP) in the presence of a divalent magnesium cofactor (Figure 1A). SmPGTs are highly selective for the nucleotide diphosphate sugar moiety, which is identifiable through the SmPGT sequence similarity network (SSN) as confirmed by large scale activity analysis (Durand et al. 2024). To further probe SmPGT protein dynamics in a bilayer system, we conducted all-atom molecular dynamics simulations prepared using the CHARMM-GUI protocol as previously described (Wu et al. 2014a; Majumder et al. 2023). We identified dynamic features by comparing the root-mean-squared fluctuations (RMSF) and the dominant motion in principal component analysis (PCA) binned by amino acid residue (Figure 1B). These dynamics were then compared to substrate-specific structural features of AlphaFold2 predictions of the SmPGT SSN. The enzyme of interest is structurally aligned to all other members of the network, and the average lDDT score per residue can be computed between a protein and the substrate-specific clusters and between the protein and the entire SmPGT network (Figure 1C).

We conducted this analysis for three representative SmPGTs that act on different UDP-sugar substrates, namely: *Cc* PglC (UDP-diNAcBac), *Acinetobacter baumannii (Ab)* ItrA4 (UDP-Gal), and *Bacillus fragilis (Bf)* WcfS (UDP-fucNAc4N) (Figure 1D). All three PGTs feature a dominant first principal component in which an analogous mobile element (ranging from 20-30 residues in length) shows the highest degree of concerted motion (Figure 1E). The most dynamic features in simulation closely correlate with those that are structurally conserved, specifically within a given substrate-specific cluster (Figure 1F). This observation suggests that dynamic structural motifs of small PGTs across the superfamily evolved to better recognize distinct UDP-sugars and catalyze their transfer.

To investigate details of the UDP-sugar binding pocket we next conducted MD simulations with the cognate sugar substrate and UndP present. We used Chai-1 to generate an initial docked structure, which reliably places the Mg^2+^ and UndP consistent with the crystal structures and prior simulations (Majumder et al. 2023; Chai et al. 2024). Although the UDP-sugar binding site is unknown, a number of key residues are understood to interact with the UDP-sugar substrate (Das et al. 2017b). These residues are also predicted to interact with the diphosphate moiety and coordinate the magnesium ion in Chai-1 structures (Figure 2A-B, magenta residues). Intriguingly, Arg112, although originally proposed to interact with the uracil moiety (Ray et al. 2018), is predicted in the opposite orientation with Arg79 instead performing a similar role in uracil binding. Of particular note is the interaction of a highly conserved methionine and an aromatic residue of previously unknown function with the uracil group (M62 and F43 for *Cc* PglC) that stabilizes the bound nucleobase and anchors the position of the mobile loop. For both *Cc* PglC and *Ab* ItrA4, Chai-1 predicts a more closed protein conformation of the substrate-specific mobile loop when a UDP-sugar is present although *Bf* WcfS remains relatively closed in all Chai-1 predictions (Figure S2, supplemental videos V1-6).

**Figure 2:**
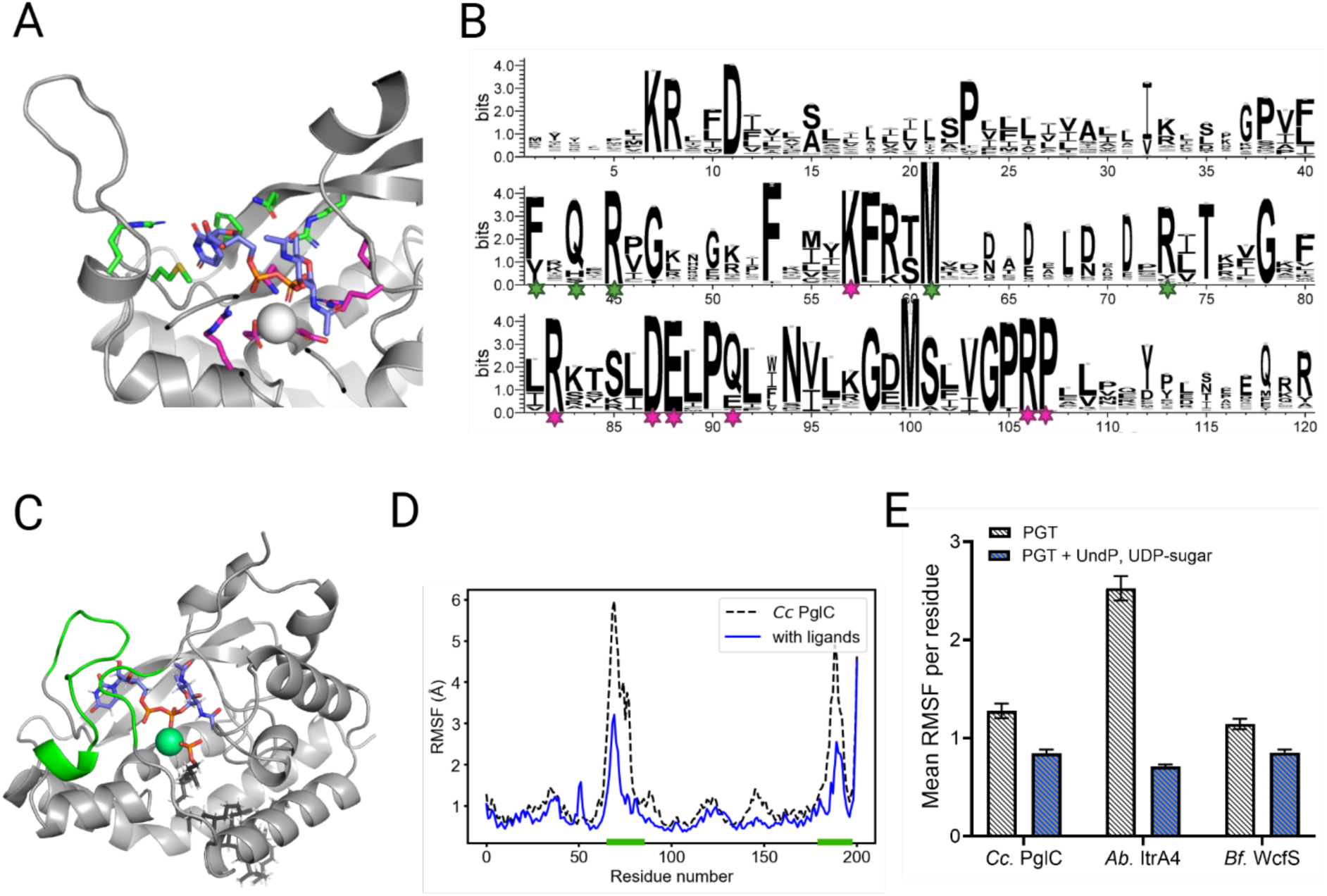
PGT stabilization with ligands. (A) Active site closeup of Chai-1 predicted pose of *Cc.* PglC with UDP-diNAcBac. Highlighted residues are conserved across the SmPGT superfamily, with magenta showing previously identified residues of the catalytic core and green showing new interactions. (B) Sequence logo plot of the SmPGT network with conserved residues from (A) highlighted. Only residues with >50% occupancy are shown to condense the scale of the plot. (C) Structure of *Cc.* PglC with UDP-diNAcBac and UndP after 500 ns simulated in a bilayer. Green highlights show significant structural rearrangement from the initial Chai-1 prediction. (D). Average RMSF per residue for *Cc* PglC simulated in a bilayer with or without ligands. (E). Mean RMSF per residue for substrate-specific motifs with and without ligands. Simulations were initialized by Chai-1 predicted complexes for the three exemplar PGTs.

We next conducted all-atom molecular dynamics with and without UndP and UDP-sugar substrates present in a phospholipid bilayer, initialized with the corresponding Chai-1 predicted structure. In all cases, the presence of substrates in this predicted orientation significantly stabilized the most dynamic residues that we have shown to be substrate specific (Figure 2D-E, Figure S2). Of note is that *Cc* PglC still showed significant variation in the position of its mobile loop in the initial 200 ns simulation with substrates. Extending this condition to a full microsecond simulation showed a significant conformational rearrangement of this loop to close the active site around UDP-diNAcBac (Figure 2C, supplementary video V7). Taken together, these results suggest that Chai-1 is able to reasonably predict the closed conformation of *Ab* ItrA4 and *Bf* WcfS when bound to their respective UDP-sugar substrates but could not fully capture the more dramatic structural rearrangement of *Cc* PglC. Nevertheless, the Chai-1 predictions enabled us to improve hypothesized PGT binding poses and substantially reduce the search space of poses and interactions using demanding molecular dynamics simulations. Critically, the analysis also identified substrate-protein interactions that could not have been determined otherwise due to challenges with the determination of high-resolution crystal structures of the unusual monotopic protein.

### Experimental platform to observe ligand-dependent conformational changes

To investigate the predicted SmPGT dynamics in a physiologically relevant membrane context and its dependence on the presence of bound ligands, we developed a platform for labeling and immobilizing a PGT in a native-like liponanoparticle for smFRET. Here, we focused on the SmPGT from the N-linked glycosylation pathway of *Campylobacter jejuni*, *Cj* PglC. Compared to *C. concisus*, *C. jejuni* is a more pathogenic strain of *Campylobacter* responsible for the majority of infections from the genus (Costa and Iraola 2019). *Cc* and *Cj* PglC are both highly active *in vitro* and specific to UDP-diNAcBac, have 63.4% sequence identity and extremely similar AF predicted structures (RMSD = 0.527 Å) which are in turn < 1.51 Å RMSD from the experimental crystal structure of PglC from *Campylobacter concisus*. Additionally, *Cj* PglC has previously been evaluated against a library of nucleoside analogs developed as potential inhibitors (Durand et al. 2024). Therefore, *Cj* PglC is a valuable test where orthogonal approaches to determine ligand binding can better inform the design and development of inhibitory small molecules.

We established a detergent-free purification pipeline with orthogonal labeling of two specific residues with fluorescent dyes for single-molecule FRET (Figure 3). For the site-specific introduction of Cy3 and Cy5 fluorophores for FRET we deployed two efficient orthogonal conjugation strategies to minimize heterogeneity in the sample. As *Cj* PglC has no native cysteine residues, we applied site-directed mutagenesis to incorporate cysteine at various sites for thiolate targeted alkylation with a sulfoCy5 maleimide derivative. For Cy3 incorporation, we applied non-canonical amino acid mutagenesis using the pyrrolysine system (Wan et al. 2014), with an engineered tRNA synthetase/tRNA^TAG^ pair for bicyclononyne-lysine (BCN) incorporation at the amber stop-codon (TAG) (Mukai et al. 2015; Swiecicki et al. 2020; Bartoschek et al. 2021). Both Cys and BCN incorporation sites were chosen at positions of low conservation across SmPGTs. The BCN sites were further screened for suppression in the absence of BCN (Figure S3A), from which we selected residue L166 for non-canonical amino acid mutagenesis. We selected two dual-variants: one with mutagenesis sites at E68 and L166 to measure the average position of the mobile loop, and one with modification at E124 and L166, which are significantly below the Förster radius of the Cy3-Cy5 FRET pair to provide a positive control (Figure 3B). Both variants were overexpressed in a strain of *E. coli* without specific assignment to the amber stop codon, solubilized from the cell envelope fraction into styrene-maleic acid lipo-nanoparticles (SMALPs), purified with a Ni-NTA pulldown (Figure S3B), and sequentially labeled with sulfoCy5 maleimide and Cy3-tetrazine (Figure 3C). This process ensured a native-like membrane environment for *in vitro* measurements while eliminating intermediate detergent solubilization, which can be highly variable and deleterious to enzyme activity and stability (Arbour Christine et al. 2023). Dual-labeled SMALPs were subsequently run on a size exclusion column to remove excess dye and remove any remaining SMA or off-target SMALPs (Figure S3C-D). Activity was confirmed for dual variants in SMALP using UMP-Glo™ (Figure S3E), and a radioactivity-based activity assay, which established the compatibility of *Cj* PglC in the presence of the oxygen scavenging system used to reduce fluorophore photobleaching and triplet-state quenching (Figure S3F) (Das et al. 2016; Arbour Christine et al. 2023).

**Figure 3:**
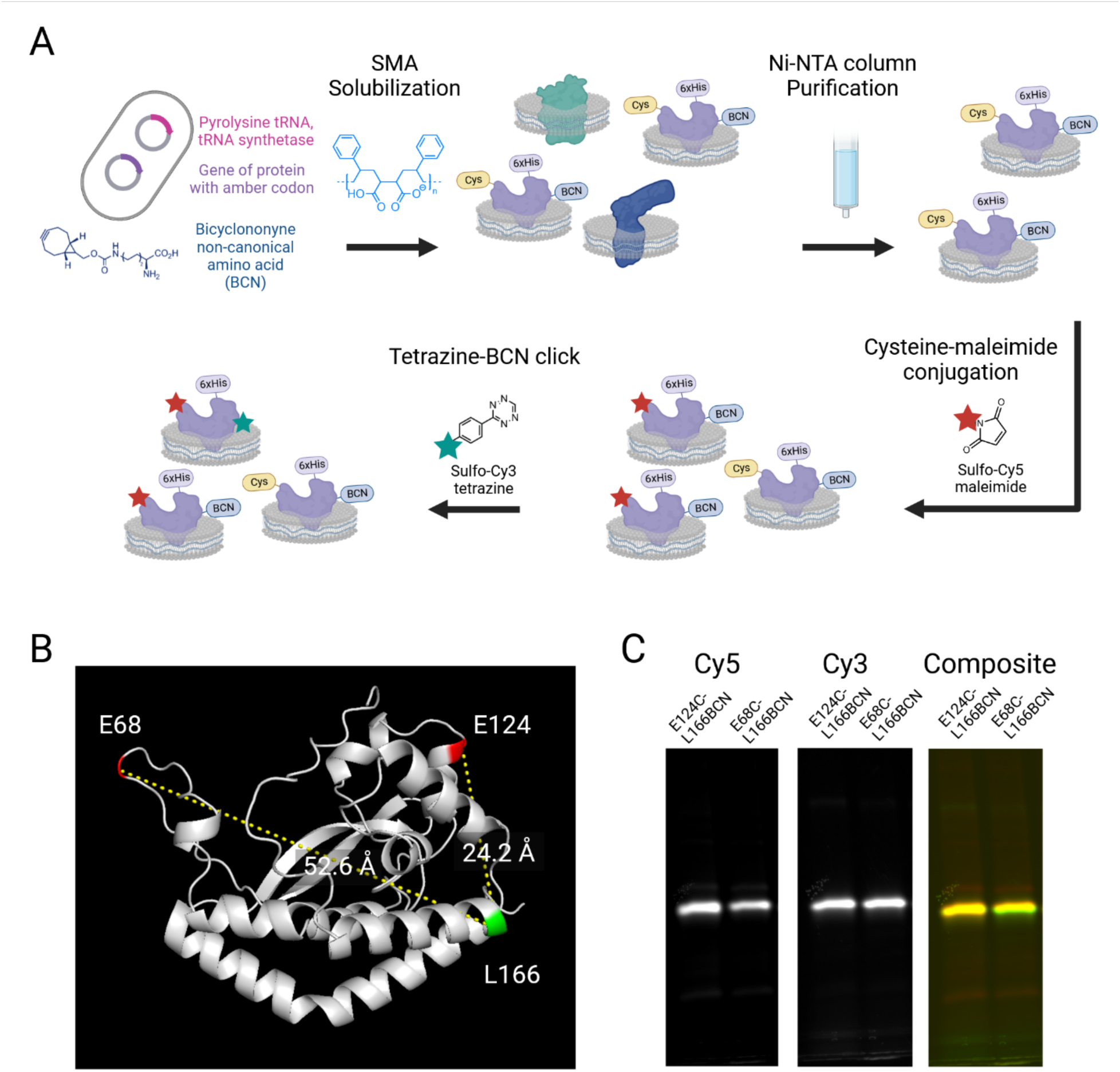
Solubilization and labeling of Cj PglC. (A) Schematic showing protein expression, solubilization, and sequential dual labeling in SMA liponanoparticles. (B) AF structure of Cj PglC showing residue sites for fluorophore conjugation in loop-labeled variant (E68C-L166BCN) and positive control (E124C-L166BCN). (C) Fluorescence gel showing both dual variants purified in SMALPs and labeled with Cy3 and Cy5 fluorophores.

For Total Internal Reflection Fluorescence (TIRF) microscopy we used a dual camera setup with a dichroic mirror splitting at 640 nm (Figure S4A). To calibrate the system for smFRET, we used Cy3 and Cy5 labeled DNA oligos tethered to PEG-biotin coated glass coverslips for camera registration and γ correction (Figure S4B-F). To tether the protein-SMALP assemblies to the glass surface we prepared PEG-modified glass coverslips with a low density of PEG-NTA instead of PEG-biotin (Figure S5A). We observed a high level of specificity for SMALPs whether a His_6_-tag or a Strep-II tag was used for protein solubilization (Figure S5B), and an excess of soluble protein with a His_6_-tag was unable to compete for these surface binding sites (Figure S5C). This suggests that the interactions between SMA in these liponanoparticles and the Ni-NTA on the surface is sufficient to reliably attach solubilized protein and show enough avidity in this geometry to supersede His_6_-tag anchoring. This illustrates Ni-NTA can be used to anchor SMALPs to glass slide surfaces directly, greatly simplifying experimental design of protein constructs.

We first compared the average FRET observed between the loop-labeled variant E68C-L166BCN with the high-FRET control variant E124C-L166BCN (Figure 4A). For each condition, videos were taken in a single batch under identical conditions. To confirm the presence of both fluorophores, we alternated excitation between donor and acceptor every frame and examined the population that showed donor emission recovery upon acceptor photobleaching while maintaining a near-constant total emission intensity (γ correction factor near unity) (Figure 4B). In this way we filtered single-molecule traces to an ensemble of approximately 50-100 traces per condition. We examined the distributions of the FRET proximity ratio (Figure 4C). The control variant showed a dominant peak at saturating values of FRET where donor emission was undetectable relative to noise until acceptor bleaching. In contrast, the loop variant featured a smaller saturating peak as well as a broader distribution of intermediate FRET values. This bimodal distribution indicates the presence of an additional state where the mobile loop is in a closed configuration, closer to the main soluble domain of the protein. To assess the relative frequency of this state in the distribution, we fit the distribution to the sum of two Gaussians (Figure 4D). Note that the control distribution is not perfectly fit by a single Gaussian distribution due to a small proportion of lower FRET traces. Therefore, we quantify the relative proportion of saturating FRET as a comparative measurement describing how well we can distinguish FRET transfer at intermediate and close distances (Figure 4E).

**Figure 4:**
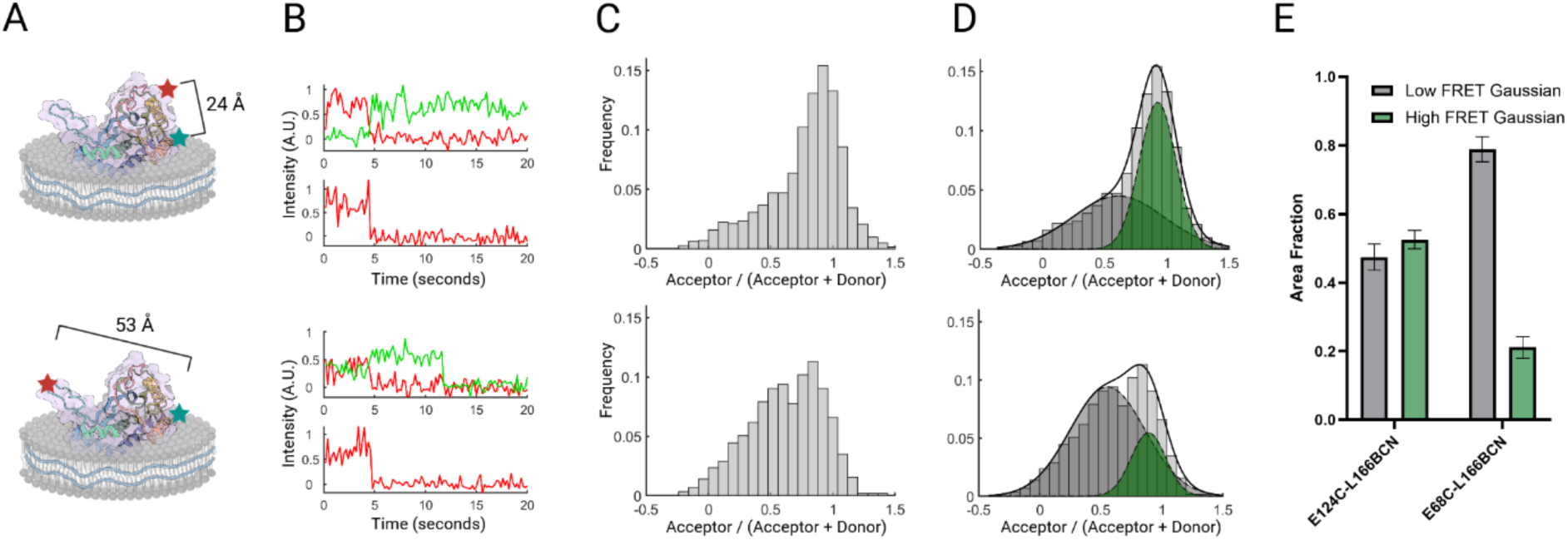
Analysis of FRET from dual-labeled *Cj* PglC variants. (A) Labeling scheme of control variant vs. loop variant with associated α-carbon to α-carbon distances. (B) Sample intensity traces. Top panel is under 532 nm illumination, where green is Cy3 emission and red is Cy5 emission. Bottom panel is 640 nm excitation, taken in alternating frames. (C) FRET distributions over a larger dataset for each variant (N = 110 and 74, respectively). (D) The kernel density estimation of each distribution from (C) is fit with two Gaussians to show relative proportion of saturating and sub-saturating FRET in the ensemble. (E) Area fraction of Gaussian fits by labeling variant. Error bars are set by the 95% confidence interval of fitting.

We proceeded to observe potential changes in the FRET distribution upon the addition of small molecule ligands. We used nucleoside mimetic compounds designed to be competitive inhibitors of PGTs and selected compounds with the same UDP mimetic backbone but with variable groups at the sugar mimetic moiety that showed a variety of inhibitory properties (Figure 5A-B) (Arbour and Imperiali 2022). The best inhibitors are NA-1 and NA-4, with reported K_i_ values of 5.7 ± 1.3 μM and 9.7 ± 0.7 μM respectively based on the UMP-Glo assay and which compete for the UDP-sugar binding site (Figure S6). To evaluate the effect of each compound on closure of the mobile loop, we measured the distribution of FRET observed from a population of loop-labeled *Cj* PglC SMALPs before and after the addition of a given ligand. This was done to check for consistency of the control distribution when imaging on different days.

**Figure 5:**
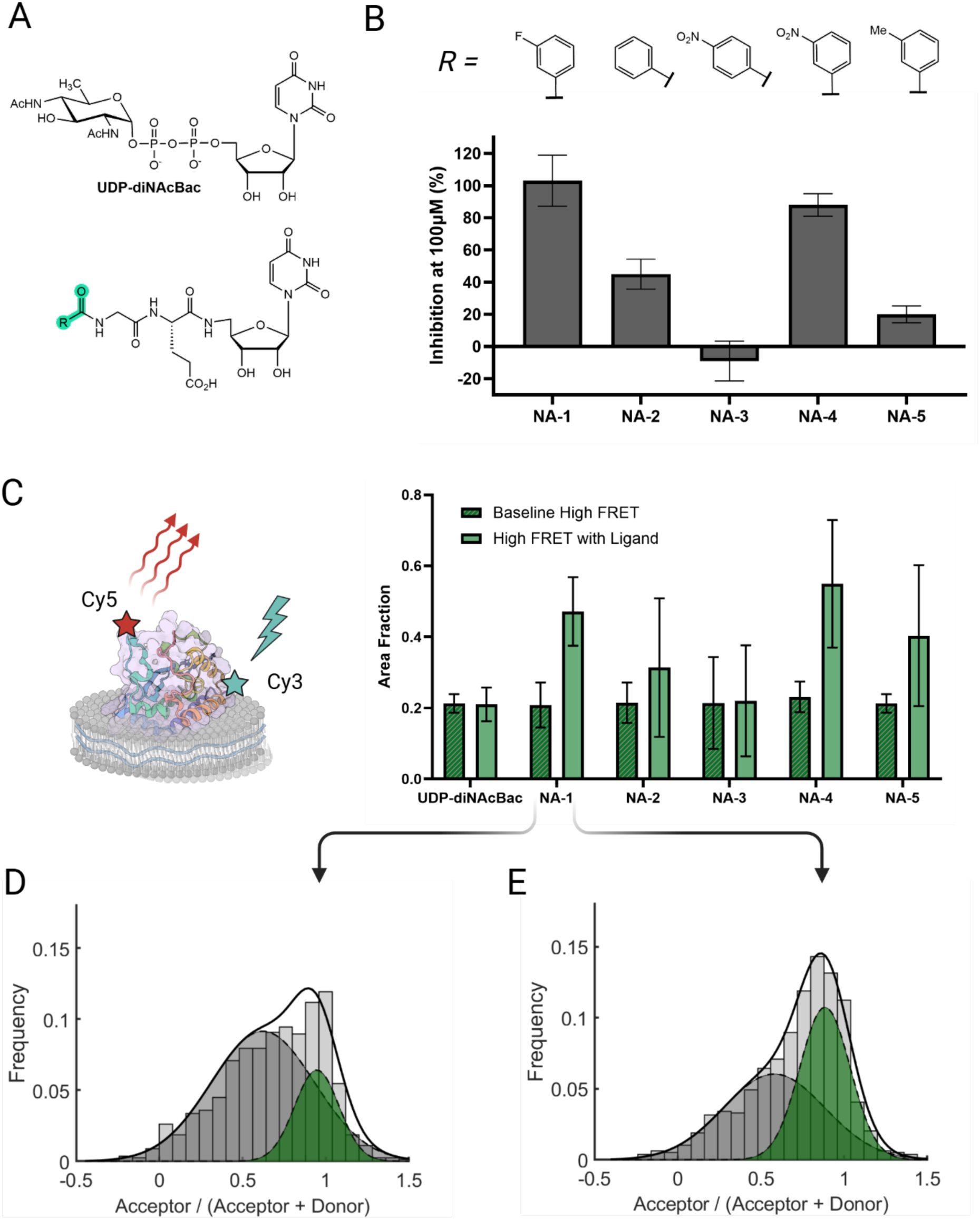
C*j* PglC mobile loop closure in the presence of synthetic nucleoside analogs. (A) <u>Top</u>: Chemical structure of UDP-diNAcBac. <u>Bottom</u>: Nucleoside analog (NA) scaffold structure. Variable R group highlighted in green. (B) <u>Top</u>: Variable R group for panel of five NA inhibitors. <u>Bottom</u>: Percent inhibition of *Cj* PglC. Activity was measured via UMP-Glo with and without ligand to calculate percent inhibition. (C) Fraction of high-FRET observed in distribution of single-molecule traces before and after addition of ligand. (D-E) <u>Left:</u> Control FRET ratio distribution and dual-Gaussian fits of loop-labeled variant before addition of NA-1. <u>Right</u> FRET ratio distribution after addition of NA-1, showing increased occurrence of high-FRET state.

In these analyses, we did not observe a significant difference in FRET when adding the cognate UDP-sugar substrate for *Cj* PglC, however, the nucleoside analogs show a significant correlation between percent inhibition and closure of the mobile loop (Figure 5C-E). This suggests that the time scale at which UDP-diNAcBac reacts to form the covalent intermediate is faster than the highest frame rate at which we can measure sufficient signal to distinguish different average loop positions (about 2.5 FPS). As a class of competitive inhibitors, the percent inhibition of the non-hydrolyzable analogs is a proxy for binding affinity to the active site. The correlation between high FRET and inhibition therefore indicates that closure of the mobile loop correlates with ligand binding, and that tighter binders will result in a higher frequency of the closed conformational state that can be observed in the distribution of smFRET traces.

## Discussion

### Integrating computational and experimental techniques informed by the membrane environment

As more is discovered about the role of local membrane composition on the dynamics of membrane proteins, characterization techniques will need to account for local bilayer chemistry and geometry (Levental and Lyman 2023). This is especially vital in the era of ubiquitous protein structure prediction and emerging advances in molecular docking and complex prediction based on large scale machine learning, where precise details of the information used are obscured within a sea of bioinformatic and structural training data. To complement top-down efforts to characterize the information used by ML structure prediction models (Zhang et al. 2024), cross-validation that explicitly accounts for the physicochemical effects of membranes on protein structure and function are vital. The methodology developed in this work demonstrates the cross-validation of ML based structural bioinformatics with molecular dynamics, biochemical characterization, and smFRET microscopy to study protein dynamics in the context of a phospholipid bilayers system. For proteins like the SmPGTs that are exceptionally challenging to characterize structurally it is critical to know what level of significance to assign different protein-ligand complex predictions.

### PGT conformational dynamics are connected to UDP-sugar binding

Our combined bioinformatic, computational, and experimental characterization of *Cj* PglC and close homologs independently correlate dynamics of the substrate-specific SmPGT mobile loop to the binding of soluble ligands and indicate a closed conformational bound state.

The Chai-1 predictions utilized in this study identified a uridine binding motif that required partial closure of the SmPGT mobile loop for repositioning amino acid side chains (namely F36, M61, and R73 in Figure 2B). The ability to predict binding poses with conformational changes is an advantage for these emerging computational methods, but proteins like PglC illustrate that more dramatic structural dynamics may necessitate all-atom molecular dynamics and smFRET measurements to identify.

Screening with synthetic nucleoside analog inhibitors allows us to demonstrate that closure of the mobile loop plays a role in binding nucleoside analogs in the UDP-sugar active site. The fact that the analogs are designed as substrate mimics that cannot be hydrolyzed allows corroboration on the role of the conformational changes on UDP-sugar binding. In this study we did not achieve sufficient sensitivity and temporal resolution to measure kinetics of the closed and opened states, but the clear correlation of high FRET with better binding competitive inhibitors corroborates the simulated dynamics of UDP-sugar binding.

This work lays the foundation for future mutagenesis studies and single-molecule kinetics to identify residues responsible for ligand specificity and further characterize the bound state, as well as further computational studies that generalize to the superfamily of SmPGTs. As more potent inhibitors are generated, they can be evaluated with this workflow to identify protein-ligand interactions.

### Modular design of competitive inhibitors for specific protein-ligand interactions

The purification, labeling, and surface-tethering platform allows us to use single-molecule FRET microscopy to visualize conformational changes in small membrane proteins *in situ*. When combined with a panel of small molecule ligands, we can also identify ligand-specific conformational changes. The SmPGTs and the library of UDP-sugar analog inhibitors are an ideal test case to study the limits of complex prediction accuracy, as these proteins are small enough to be computationally tractable for a variety of computational approaches and substrate-specific enough that individual changes in stereo- and/or regiochemistry can dramatically affect binding. Additionally, substrate-mimetic nucleoside analog inhibitors allow direct comparison to UDP-sugar binding, and the modular design streamlines force-field optimization for further MD studies. Specific interactions may then be independently tested for binding and the induction of a conformational change by comparing nucleoside analog chemistry with our pipeline and targeted mutagenesis studies with data from the smFRET platform. Ligands can be screened for inhibition and induced fit by smFRET in to streamline selection for structural studies to capture bound conformations.

## Materials and Methods

### Protein structure prediction

AlphaFold 2 structures were predicted using LocalColabFold v1.5.5 (Mirdita et al. 2022) run with flags –amber, --templates, --num-recycle 3, --use-gpu-relax on an Nvidia GEFORCE RTX 2080 GPU. Chai-1 v0.4.0 predictions were run locally using an Nvidia GEFORCE RTX 3070 GPU using the Colabfold MSA server with default parameters.

### All-atom MD simulations

All proteins were simulated in a lipid bilayer composed of 67 mol% 1-palmitoyl-2-oleoyl-sn-glycero-3-phosphoethanolamine (POPE), 23 mol% 1-palmitoyl-2-oleoyl-sn-glycero-3-phosphoglycerol (POPG), and 10 mol% cardiolipin (CL) of defined acyl-chain composition using the CHARMM36 all-atom force field (MacKerell et al. 1998). Simulations were prepared using the CHARMM-GUI protocol and the CgenFF to generate topologies for UndP and UDP-sugars (Vanommeslaeghe et al. 2010; Wu et al. 2014b). Analyses were conducted using the MDAnalysis python package (Michaud-Agrawal et al. 2011; 2016).

### PGT structural analysis

AlphaFold2 structure predictions of the PGT sequence similarity network (SSN) were aligned and analyzed using analysis tools from the Foldseek repository. Example PGT structures were queried against the entire PGT network and aligned using Tmalign, and the per-residue lDDT score was calculated. The average values were calculated in both the network and the substrate-specific clusters to identify substrate-specific structures.

### Protein expression

Wild-type and single-cysteine variants of PglC were expressed with a C-terminal His_6_-tag or with a Strep-Tag and *in vivo* SUMO tag cleavage as previously described (Entova et al. 2018; Dodge et al. 2023). For BCN incorporation, a culture of *E. coli* B.95 cells (B.95.ΔA) transformed with the pEvol PylRS AF and pET24 plasmids containing *Cj* PglC amber codon variants and grown as described in the Supporting Methods (Swiecicki et al. 2020).

PglC variants were purified in SMA200 liponanoparticles as previously described, albeit with a modified washing and labeling procedure on resin to reduce contamination due to non-specific sticking of SMALPs sticking to the resin and excess dye to SMALPs. Purified protein was further cleaned up by size exclusion chromatography to remove excess dye and SMA. Protein concentration was quantified with a Pierce™ BCA protein assay using WT SUMO-PglC as a standard, and fluorophore concentration was determined by UV-vis to calculate labeling efficiency.

### Activity and Inhibition Assays

The activity of SMALP solubilized PglC variants was measured using the UMP-Glo luminescence assay as previously described with modifications to the assay buffer for compatibility with liponanoparticles (Figure S3A) (Entova et al. 2018).

Inhibition assays were conducted on detergent solubilized SUMO-PglC by measuring activity by UMP-Glo with or without 100 μM of a given nucleoside analog. For the two highest inhibition compounds, PglC conversion was measured as a function of inhibitor concentration to calculate the IC_50_ values. The different compound concentrations were then assayed with a range of substrate concentrations to procure the inhibitory constants (K_i_). The compounds were determined to be competitive inhibitors as demonstrated by the changing slopes, in addition to the lines converging upon the y-axis in the Lineweaver-Burk plots (Figure S6).

Additional activity assays were performed using radiolabeled UDP-sugar substrate to confirm PglC PGT activity is not affected by imaging buffer with the presence of the oxygen scavenging system (Figure S3B) (Arbour Christine et al. 2023).

### Functionalized slides

Functionalized glass slides were created with mPEG-silane-2000 and either biotin-mPEG-silane-3400 (Laysan Bio) or NTA-mPEG-silane-3400 custom ordered from Nanocs. The glass cleaning and functionalization procedure was modified from the literature protocol to include a chelation step to remove divalent cation contamination (Figure S4) (Gupta et al. 2021).

### TIRF Microscopy

Fluorescence measurements were performed on an epifluorescence dual-camera TIRF microscope. DNA mapping oligo controls were ordered from Millipore Sigma and based on designs from the literature (McCann et al. 2010; Gupta et al. 2021).

Image analysis was performed using custom code written with the MATLAB image processing and curve fitting toolboxes, which is further detailed in the Supporting Methods.

## Supporting information

CcPglC

CcPglC-UDPdiNAcBac

AbItr4

AbItr4-UDP-Gal

BfWcfS

BfWcfS-UDPFucNAc4N

CcPglC-oneMicrsecond

## Acknowledgements

We thank Dr. Greg Dodge and Professor Soumi Ghosh for advice and assistance with protein expression and purification, as well as providing edits to the manuscript. We thank Theo Durand for his curation of the PGT SSN and Professor Karen Allen for detailed discussions and structural insight. We also thank Dr. Ayan Majumder and Professor John Straub for advice for all-atom MD simulation preparation and analysis.

The microscope system we used was built by the MIT Bell Lab with Dr. Charles Felts, and we thank Drs. Annie Zhang, David Driscoll, and Larry Friedman for their advice for functionalizing and testing glass coverslips for single-molecule tethering.

Figures were created with Biorender.com. This work was funded by NIH grants R01 GM131627 (to BI) and F32 GM136023 (to CAA).

## Supporting Information

### SUPPLEMENTARY TABLES

**Table S1:** Bacterial strains.

|  |  |
| --- | --- |
| DH5α | Maintained in our lab |
| BL21-DE3 RIL | Lajoie et al., 2013 [1] |
| B.95 ΔA | Mukai et al., 2015 [2] |
| C43 pAM174 | Sjodt et al., 2018 [3] |

**Table S2:**
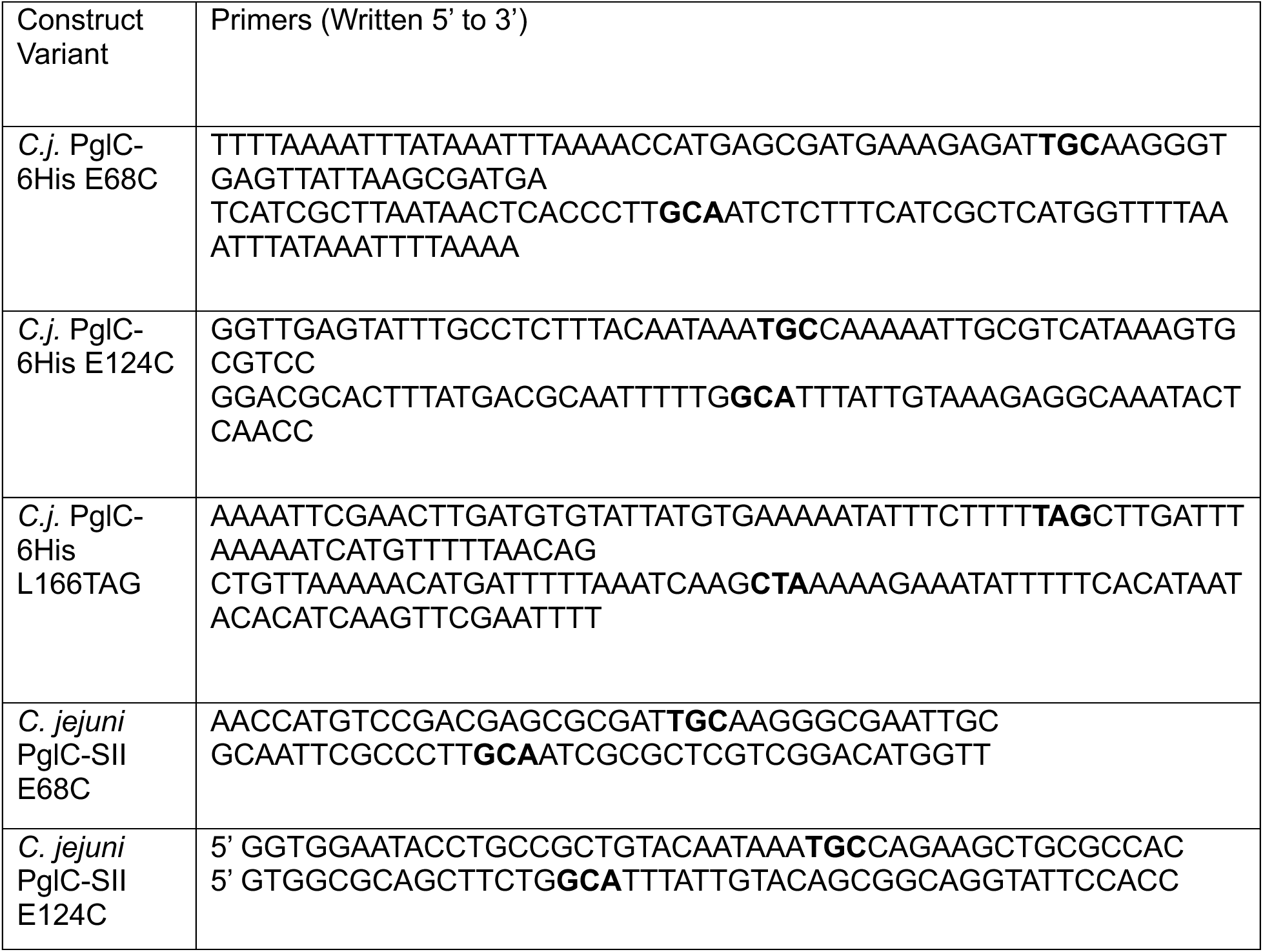
Mutagenesis primers designed with the Agilent QuikChange Primer Design tool. Mutagenesis was confirmed via Sanger sequencing.

**Table S3:**
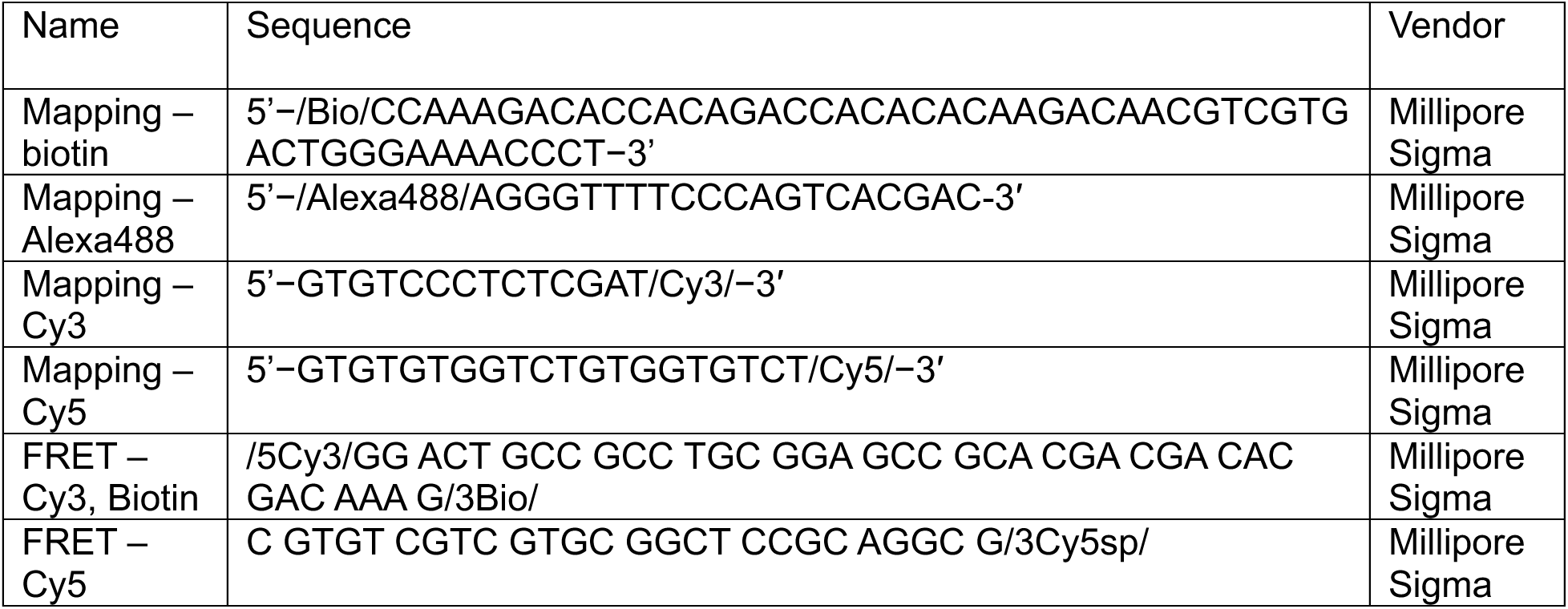
Functionalized DNA Oligos for imaging controls.

**Table S4:**
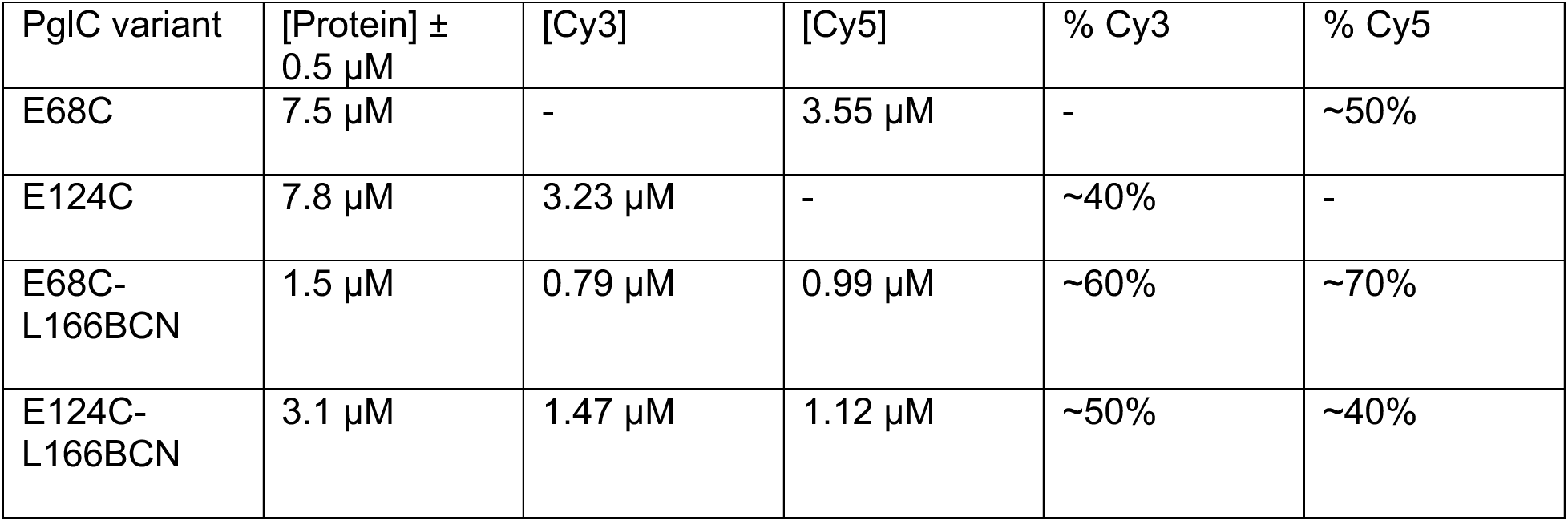
Protein labeling quantification.

### SUPPLEMENTARY FIGURES

**Figure S1:**
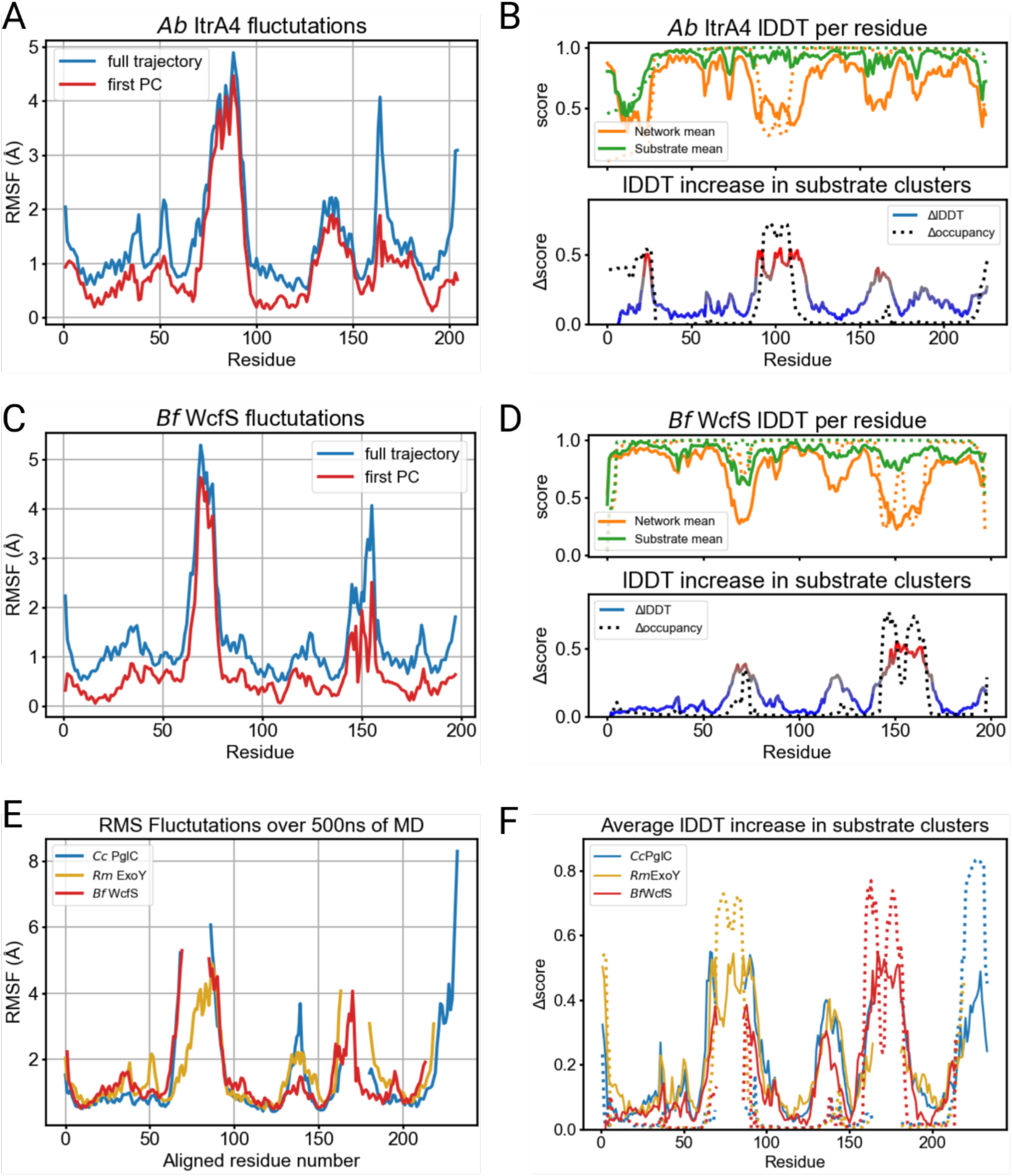
Per-residue RMSF and lDDT analysis for other PGTs. (A), (C): All fluctuations (blue) and PC1 motion (red) (B), (D): Top: lDDT per residue against all SmPGTs (orange) and against all members of the same substrate-specific cluster (green). Solid lines indicate average lDDT against residues aligned to the same position and dashed lines indicate fractional occupancy of aligned residue at that site. (E): RMS fluctuations for all three PGTs when aligned to each other by amino acid sequence. Analogous regions of each protein show elevated dynamics. (F): Difference between network and cluster lDDT aligned by sequence. The same analogous and dynamic structural motifs are highly substrate specific. Relative occupancy only highlights substrate-specific regions of variable length.

**Figure S2:**
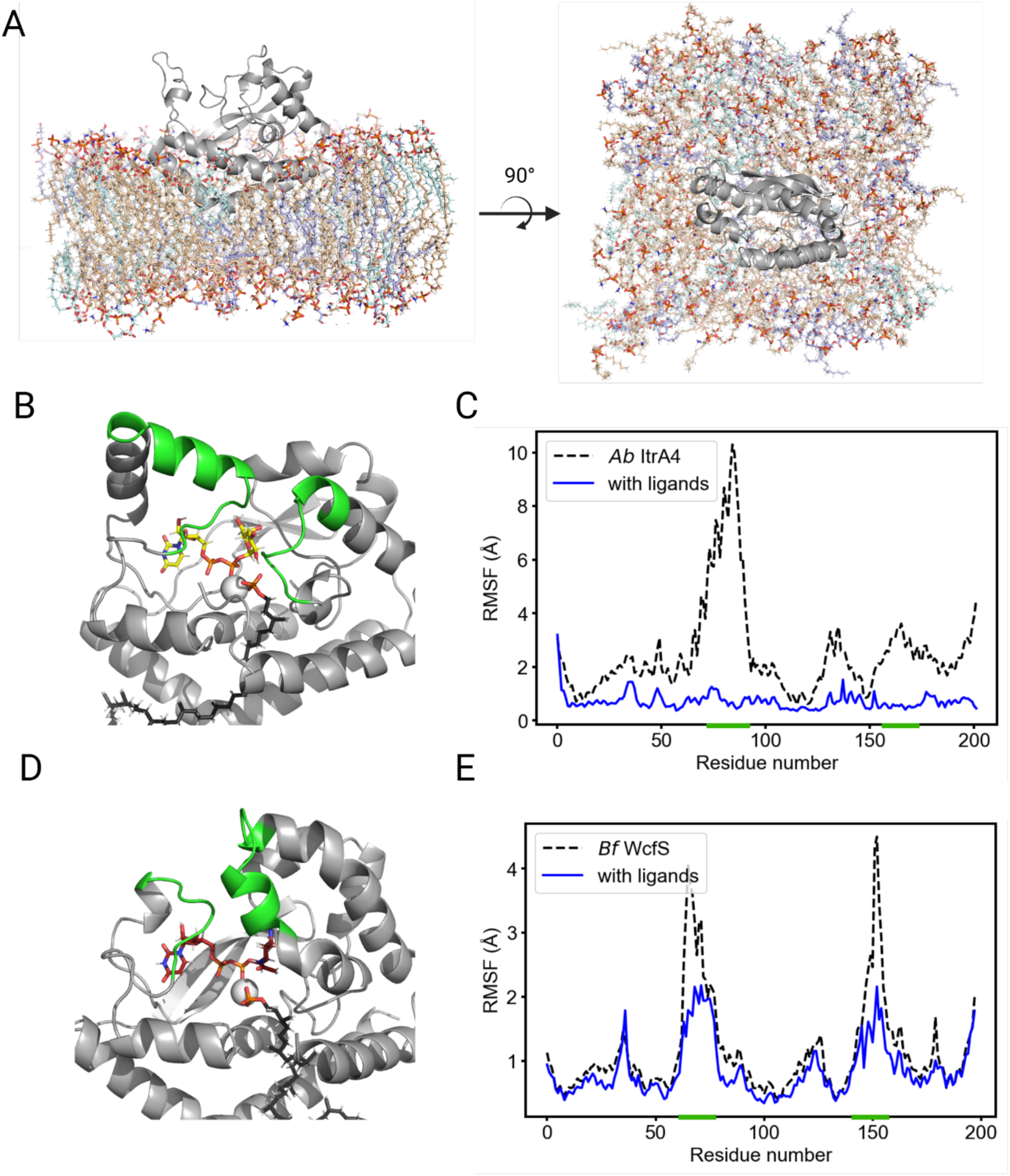
**Protein stabilization in all-atom MD**. (A): Snapshot of *C. concisus* PglC equilibrated in membrane. Tan is POPE, blue is POPG, and cyan is Cardiolipin. (B): Snapshot of *A. baumannii*. ItrA4 with UDP-galactose (yellow) and UndP (black) after 200 ns of simulation in a phospholipid bilayer. Green highlighted regions showed the most dramatic stabilization and correspond to substrate-specific structures. (C): Average RMSF per residue over 200 ns of simulation with or without ligands. Green highlighted regions shown on the x-axis. (D): *B. fragilis*. WcfS snapshot with UDP-fucNAc4N (red) and UndP (black) after 200 ns simulation. (E) WcfS fluctuations per residue with or without ligands present.

**Figure S3:**
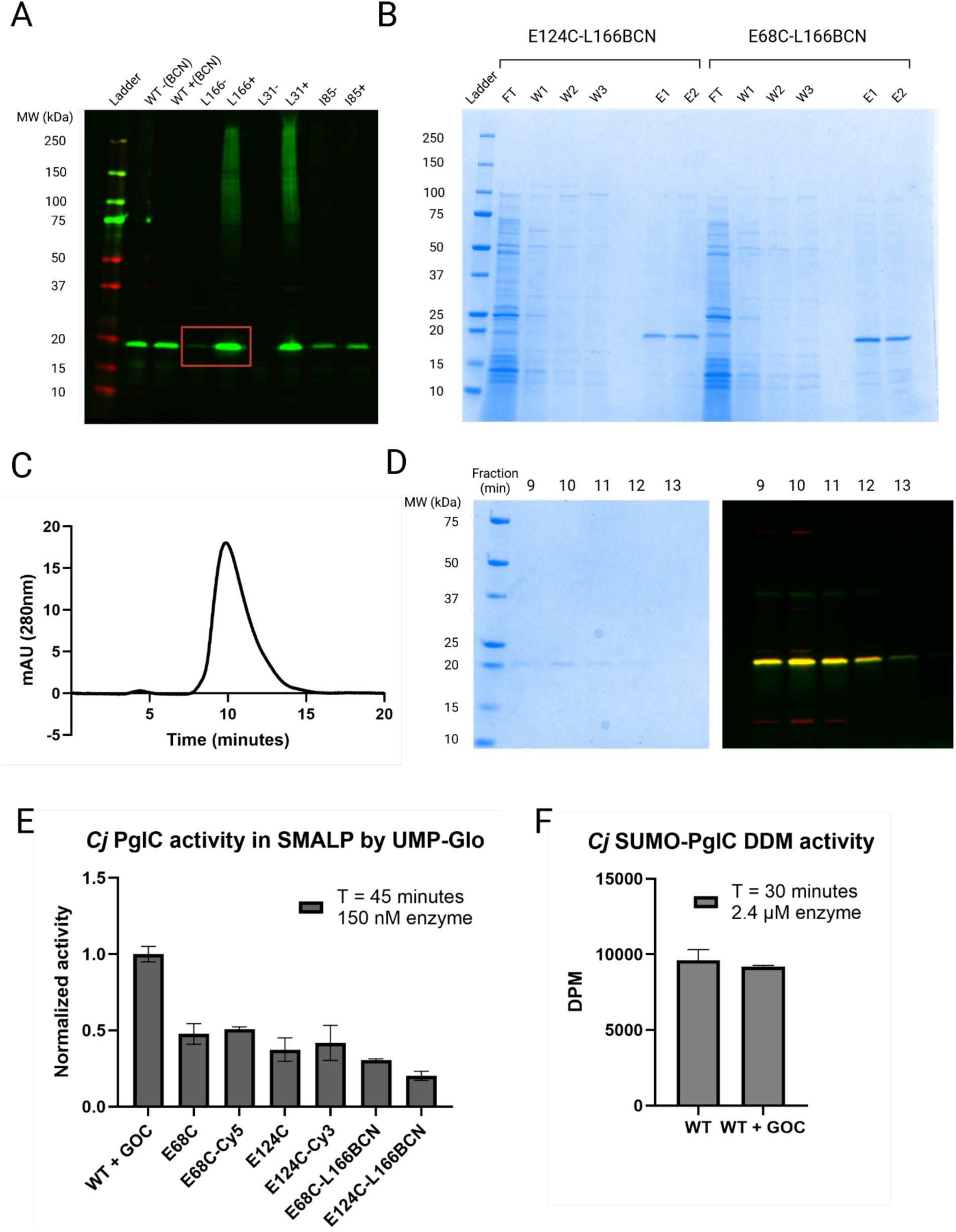
Purification and characterization details for *Cj.* PglC dual-labeled variants. (A): Fluorescence western of test cultures of B95 cells expressing BCN point variants of Cj.PglC. L166 showed strong suppression without BCN and the best expression with BCN present. (B). Coomassie gel of initial Ni-NTA purification of BCN, cysteine dual variants. (C): UV absorption trace from size exclusion column on loop-labeled variant E68C-L166BCN. 0.5 ml fractions were collected every 30 seconds. (D) Left: Coomassie gel of every-other SEC fraction, labeled by minute collected. Right: Fluorescence composite of SEC fractions taken prior to Coomassie staining. Fractions were combined for the fluorescence gel in Figure 3. (E): Activity of Cj PglC variants in the UMP-Glo assay. (F): Radioactivity-based assay of detergent-solubilized SUMO-PglC. Oxygen scavenging reaction does not significantly affect activity.

**Figure S4:**
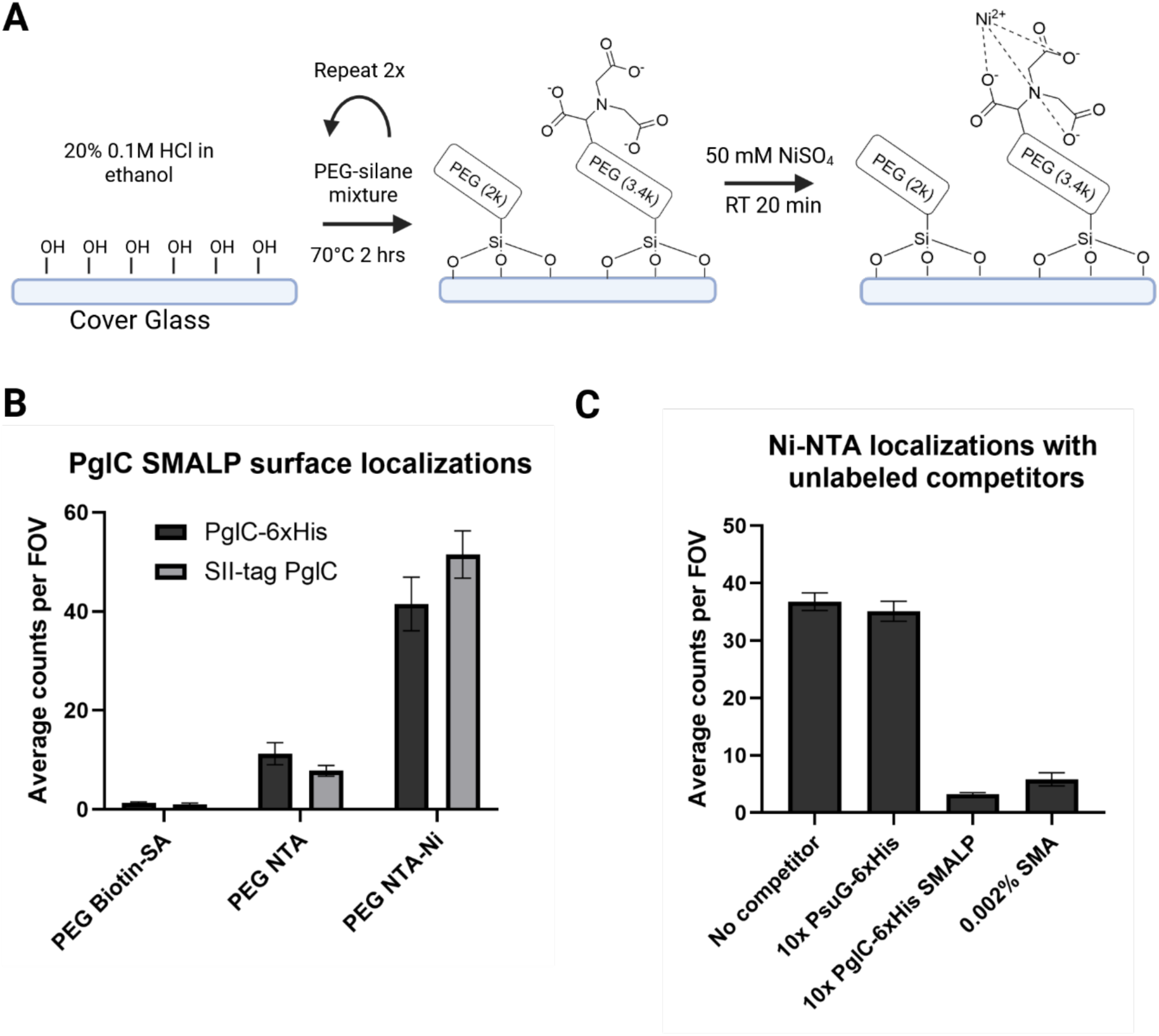
Slide synthesis and specificity controls. (A) Surface functionalization scheme for labeling protein. (B) Number of single-molecule localizations (determined by single-step photobleaching) per field of view for different slide surface chemistries, 50:1 mass ratio of PEG to functionalized-PEG. Similar behavior observed for SMALPs purified with 6xHis and SII tags. (C) Competition experiments to test for specificity of surface binding. Unlabeled SMALPs or free SMA compete with labeled SMALPs for surface binding sites, while soluble protein with a 6xHis tag does not. All error bars are given as SEM across 10 – 20 separate fields of view (FOV).

**Figure S5:**
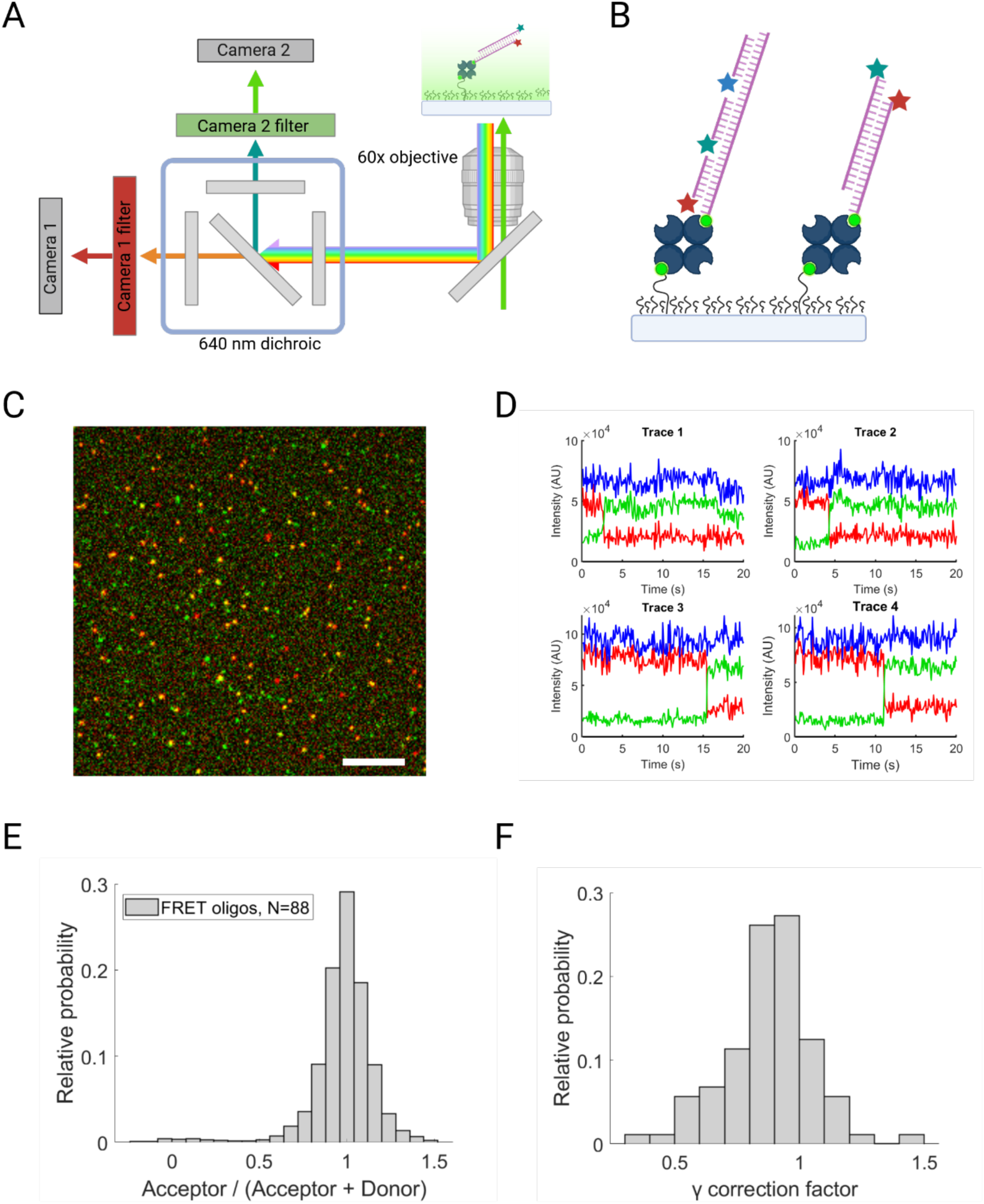
Microscope setup and DNA-oligo controls. (A): Microscope diagram illustrating dual camera dichroic setup. (B): Schematic of biotin-streptavidin slides and mapping oligomers (left) versus FRET oligos (right). (C) Overlay of Cy3 and Cy5 from mapping oligomers illustrating manual camera registration. One pixel tolerance sufficed to correctly map colocalized points across the field of view. (D) Sample FRET traces illustrating γ factor near 1 after manual adjusting gain between cameras. Green is Cy3 emission, red is Cy5 emission, and blue is total intensity that will remain constant for a γ factor of 1. (E): FRET ratio distribution for DNA FRET oligos. (F): Corresponding γ distribution for the same FRET traces of these oligos. For subsequent experiments γ outside of [0.75,1.25] was used to filter lower quality traces. 9

**Figure S6:**
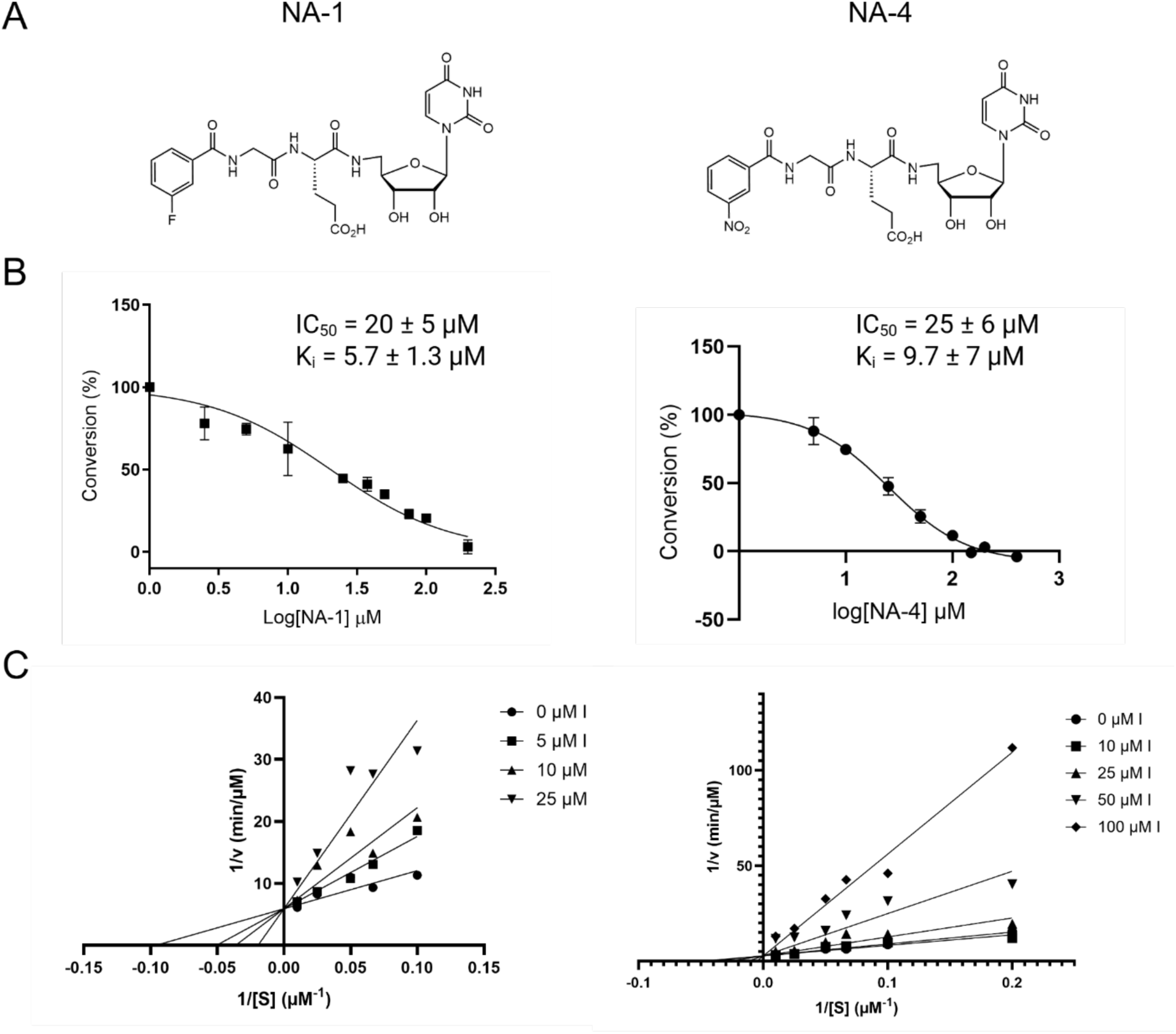
Kinetic **analysis of strongest nucleoside inhibitors NA-1 and NA-4.** (A): Chemical structures of nucleoside analog inhibitors. (B): IC50 curves of select inhibitors with C. jejuni PglC, measured by UMP-Glo monitoring luminescence. Percent conversion was determined after pre-incubation with inhibitor as described by the UMP-Glo Assay procedure, in reference to control with no inhibitor. Error bars indicate mean and error ± SD; n = 2. (C): Lineweaver-Burk plot of C. jejuni PglC with compounds NA-1 and NA-4 and UDP-diNAcBac. The substrate conversion was monitored by UMP-Glo luminescence assay.

**Figure S7:**
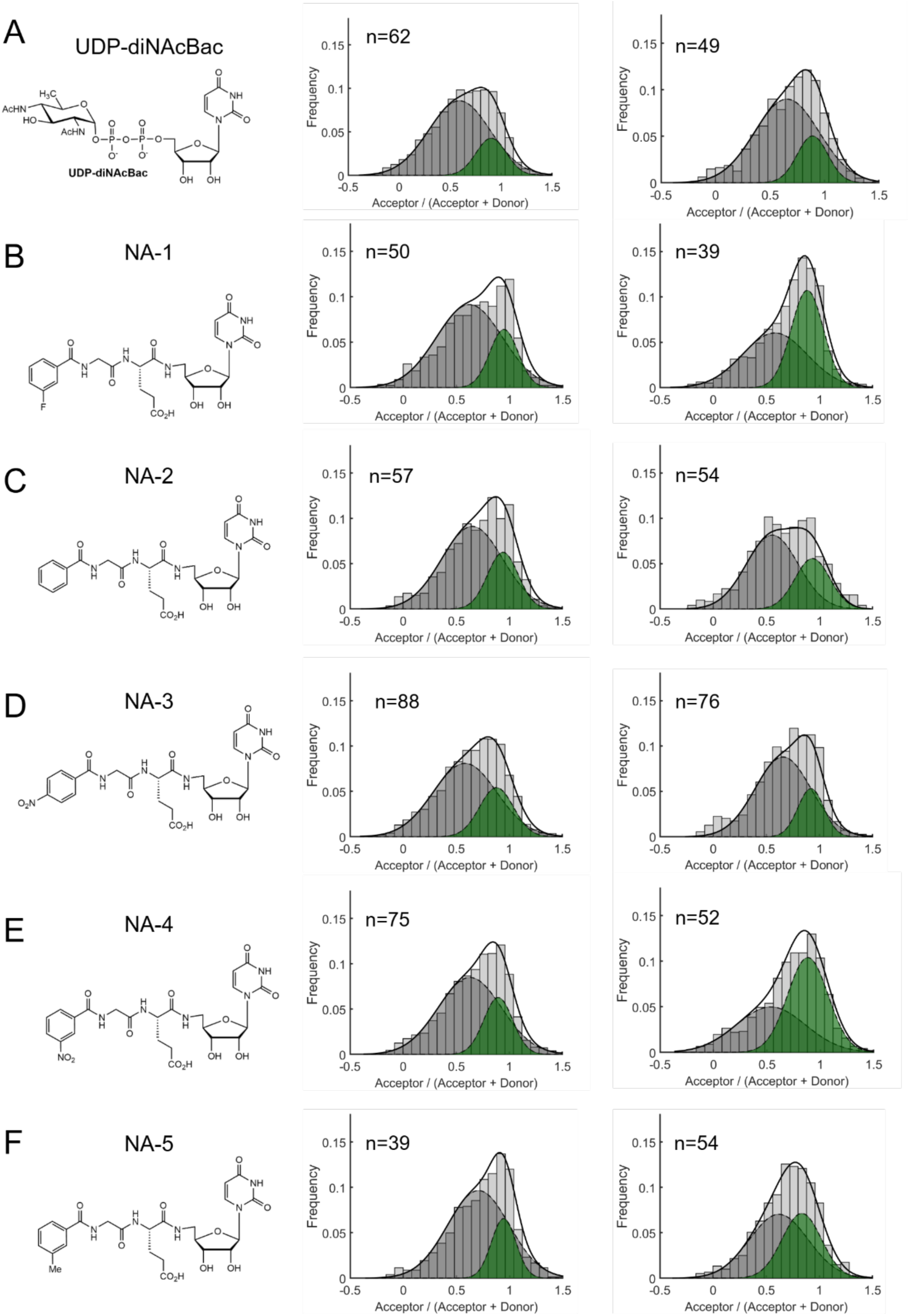
FRET ratio distributions for full ligand screen. (A)-(F): <u>Left:</u> Full chemical structure of UDP-diNAcBac or nucleoside analog. <u>Middle</u>: Control distribution and fits taken for each inhibitor without ligand present. <u>Right:</u> Distribution and fits of FRET ratios in the presence of each ligand.

### SUPPLEMENTARY METHODS

#### All-atom MD simulations

All proteins were simulated in a lipid bilayer composed of 67 mol% 1-palmitoyl-2-oleoyl-sn-glycero-3-phosphoethanolamine (POPE), 23 mol% 1-palmitoyl-2-oleoyl-sn-glycero-3-phosphoglycerol (POPG), and 10 mol% cardiolipin (CL) of defined acyl-chain composition using the CHARMM36 all-atom force field [4]. Lipid bilayer systems were solvated using the TIP3P water model with 0.15 M KCl and run at a temperature of 303 K. The CHARMM general Force-Field (CgenFF) webserver was used to generate topologies for UndP and UDP-diNAcBac [5].

Each system was equilibrated for at least 50 ns following the CHARMM-GUI protocol with additional CHARMM relaxation and inputs generated for GROMACS [6]. All simulations were performed using the GROMACS program version 2024.1 unless otherwise stated.

Principal component analysis, residue contacts, and root-mean square fluctuations were computed using the MDAnalysis library in python 3.12.

#### Structure prediction analysis

The three representative PGTs were chosen had distinct UDP-sugar substrate selectivity as confirmed via UMP-Glo [7]. Additionally, *Cc.* PglC is the only SmPGT with a solved structure at time of writing, and *Ab.* ItrA4 is a close structural and sequence analog of the catalytic domain of *Se.* WbaP which has been solved by Cryo-EM [8].

We created a local Foldseek database of the AlphaFold 2 predictions of the entire SmPGT network and queried each representative using Foldseek easy-search with flags –alignment-type 1, --exhaustive-search, --format-output “query target alntmscore lddtfull”, which performed structural alignments using the accelerated TMalign for a global alignment. The lDDT and occupancy was computed per residue from the full output by averaging over the appropriate subset of PGTs for a given cluster/network comparison.

#### Protein expression of wild-type and single-cysteine variants

Mutagenesis sites were selected at positions with low amino acid conservation scores when aligned to the family of small PGTs in the SmPGT SSN. [9] PglC from *C. jejuni* was mutated using QuikChange to replace specific residues with cysteines or the amber stop codon TAG. Primers were designed with the QuikChange online primer tool (Agilent) and successful mutations were confirmed via Sanger sequencing (Table S2).

Wild-type (WT) and SUMO-WT protein with C-terminal 6xHis purification tag were overexpressed using the Studier autoinduction method in the *E. coli* strain Bl21 [10]

For Strep-tagged variants, C43 cells harboring pAM174 [3] were transformed with a plasmid encoding *C. jejuni* PglC with an N-terminal SUMO tag – linker – dual-strep tag sequence.[11] Cells were also expressed by autoinduction with the addition of 1 g solid (L)-arabinose at OD_600_ ∼1.5 to induce SUMO cleavage before the temperature was reduced to 17 °C. Cells were harvested by centrifugation, flash frozen, and stored at - 80°C. [12]

#### Non-canonical amino acid mutagenesis

BCN variants were grown in 0.5ml test cultures (see below) and assessed by anti-His western blot for expression in the presence of BCN and suppression in its absence to ensure proper ncAA incorporation (Figure S3A).

A culture of *E. coli* B.95 cells (B.95.ΔA) transformed with the pEvol PylRS AF and pET24 plasmids containing PglC amber codon variants were incubated overnight in LB containing chloramphenicol and kanamycin (200 rpm; 37 °C). 100 mL of an LB broth shaking culture (200 rpm; 37 °C) supplemented with chloramphenicol and ampicillin for selection were inoculated with 250 µL of the overnight culture (dilution x200). A stock of 80 mM BCN unnatural amino acid in 0.2 M NaOH, 15% DMSO, was prepared. At an OD of 0.2, the stock solution of BCN unnatural amino acid was diluted down to 16 mM in 1 M HEPES buffer, pH 7.5, and finally added to final concentration of 1 mM. Simultaneously, cells were induced with 0.05 % L-arabinose (20 % stock in water). The culture was further incubated for 40 minutes at 37 ° C. Cells were finally induced with IPTG (final concentration of 0.2 mM) and incubated at 25 °C overnight. Cells were harvested by centrifugation, flash frozen, and stored at –80 °C. [12]

#### Protein purification and fluorophore labeling

WT-SUMO *Cj* PglC was purified in n-Dodecyl-β-D-Maltoside (DDM) as previously described [13].

Protein was purified in SMALP200 liponanoparticles as described in [14]. After Ni-NTA or Strep-Tactin XT 4flow resin pulldown, protein was desalted with Zeba spin desalting columns into SMALP buffer (150 mM NaCl with 50 mM HEPES at pH 8.0). Protein concentration was quantified with a Pierce™ BCA protein assay kits and incubated for 2 hours at room temperature with a 10x molar excess of sulfo-Cy3 or sulfo-Cy5 maleimide dye (Lumiprobe). For dual labeled variants, an additional labeling step was conducted for 2 hours at room temperature with Cy3-tetrazine dye.

To remove excess dye and SMA, we purified the sample using size exclusion chromatography as a final step (Figure S3C-D) [8].

#### Activity and Inhibition Assays

The activity of SMALP solubilized proteins was confirmed using the UMP-Glo luminescence assay as previously described [15]. Given the reduced turnover of UndP in SMALPs, enzyme was pre-incubated with UndP for 30 minutes and a higher concentration was used relative to previously reported assays. Assays were incubated and in SMALP buffer with 1 mM MgCl2, 10% DMSO, 100 μM UndP, and the reaction was initiated with the addition of 20 μM UDP-diNAcBac.

Inhibition assays were conducted using UMP-Glo [Promega cat. VA1130] as previously described [7], which detects UMP release from the phosphoglycosyl transferase reactions. Due to the structural similarity between the synthesized inhibitors and UMP, we also controlled for off-target inhibition of the Glo reagent enzymes by the nucleoside analogues. To correct for off-target inhibition, 1.5 µM UMP (the amount typically released during the PglC reactions) was combined with inhibitor or DMSO (control) followed by the addition of the detection reagent. The percentage of UMP signal lost was used to adjust the luminescence readout [16]. The quenching solution was prepared as described by Promega. Assays contained 10% DMSO, 0.1% Triton X-100, 0.05 mg/mL (0.76 µM) Bovine Serum Albumin (BSA), 50 mM HEPES at pH 7.5, 100 mM NaCl, 5 mM MgCl2, 20 μM UndP (from either 10X stock: 200 µM or 20X stock: 400 µM in DMSO), 20 μM UDP-diNAcBac (from 10X stock: 200 µM in Milli-Q H2O), and 0.5 nM *C. jejuni* PglC (from 10X stock: 5 nM in *Cj* PglC buffer) in a final volume of 12 μL. Inhibitors were added from a 2 mM stock in DMSO, at a final concentration of 100 μM. PglC was preincubated in the reaction mixture lacking UDP-diNAcBac for 10 min at ambient temperature. The reaction rate was predetermined to be linear over 15 min at the given concentrations. Upon the addition of UDP-diNAcBac, the reaction was allowed to proceed for 9 min before the addition of quenching solution (12 µL). The reaction mixture (20 µL) was transferred to a 96-well plate (white, nonbinding surface, Corning). The plate was shaken at low speed for 16 min and incubated for 44 min at ambient temperature, and luminescence was read using a Synergy H1 hybrid plate reader (Biotek). Data were plotted using GraphPad Prism, as percentage inhibition compared to the positive control (no inhibitor). PglC buffer for enzyme dilutions: 50 mM HEPES at pH 7.5, 100 mM NaCl, 5 mM MgCl2 with 0.2% DDM.

To establish the competitive mechanism of action of the inhibitors, the compounds were screened at varying concentrations with a range of substrate concentrations (UDP-diNAcBac 10, 15, 20, 40, and 100 µM for compound NA-1 and 5, 10, 15, 20, 40, and 100 µM for NA-4). Assays in the absence of inhibitors were performed to establish linear kinetics for *Cj* PglC (0.5 nM) with the range of substrate concentrations using modified conditions described above. The assays were then performed in the presence of inhibitors and were quenched within the linear range (6-10 min depending on substrate concentration). The reciprocal of the initial reaction velocity (1/v) was plotted against the reciprocal of the substrate concentration (1/[UDP-diNAcBac]). Both inhibitors were determined to be competitive due to the converging lines on the y-axis in the Lineweaver-Burk plots.

Additional activity assays were performed using tritiated UDP-sugar substrate to monitor product formation [17] in detergent solubilized SUMO-PglC with or without the presence of the oxygen scavenging system. Activity is reported as the disintegrations per minute (DPM) in the organic layer, with error bars given by standard deviation for n = 2 replicates.

#### Glass coverslip functionalization

Functionalized glass coverslips were created as described with biotin-mPEG-silane-3400 (Laysan Bio) or NTA-mPEG-silane-3400 custom ordered from Nanocs [18].

Coverslips were incubated on the day of the experiment with 100 mM EDTA pH 7.5 for 15 minutes to remove divalent cation contamination before washing with 10 mM HEPES pH 8. This was found to reduce non-specific tethering of SMALPs to both biotinylated or Ni-NTA slides (Figure S3B). Biotin slides were incubated with a 400x dilution of 5 mg/mL streptavidin aliquots for 1 minute before biotinylated DNA constructs were flown in and incubated for an additional 1 minute. NTA slides were incubated with 100 mM EDTA pH 8 for 15 minutes to remove divalent cation contamination, and prior to imaging, slides were then washed with 10 mM HEPES pH 8 and incubated with 50 mM NiSO4 for 30 minutes.

#### TIRF Microscopy

Fluorescence measurements were performed on an epifluorescence TIRF microscope. The setup included an Eclipse Ti microscope (Nikon) equipped with a 60× Apo-TIRF oil immersion objective lens (NA 1.49; Nikon) placed on a vibration cancellation table (TMC). Labeled proteins in the evanescent field were excited at 532 nm and fluorescence was imaged on a dual-camera ImagEM EM-CCD setup (Hamamatsu), where acceptor and donor emission were separated through a dichroic beam splitter.

DNA mapping oligo controls [19] were used to manually register the two cameras, while DNA oligos designed for high FRET transfer were used to adjust camera gain for a gamma factor approximately equal to 1 between the two color channels (Figure S5A).

The imaging buffer consists of 0.2 μm filtered SMALP buffer with 5 mg/mL bovine serum albumin and 0.05% tween 20 detergent to reduce non-specific interactions with the coverslip.

The oxygen scavenging system consisted of glucose oxidase and bovine catalase (GOC) with Trolox to reduce triplet-state quenching [18]. On the day of the experiment lyophilized aliquots of GOC were resuspended to a 10X stock solution and syringe filtered through a 0.2 μm PES membrane. Immediately prior to imaging 100 μL of oxygen scavenging buffer was mixed to a final concentration of 1X GOC (40 U/mL glucose oxidase, 1500 U/mL catalase), 1 mM MgCl2, 2 mM Trolox, 2% DMSO, 50 mM D-glucose, and 100 μM inhibitor. Trolox was diluted 100X from a pure DMSO stock, and inhibitors were diluted 10X from a 10% DMSO stock.

Protein SMALP samples were diluted to ∼10 nM in imaging buffer. Flow chambers on functionalized slides were wetted with the same buffer before labeled protein was perfused in and incubated for 30-minutes at room temperature on the slide. After washing with 200 μL of imaging buffer to remove untethered protein, the oxygen scavenging system was mixed and perfused over the slide immediately prior to imaging. Oxygen scavenger was replenished every 30 minutes to prevent buffer acidification.

#### Microscopy data analysis

Raw TIFF image stacks were background subtracted using a top-hat filter with a 4-pixel radius disk as a structuring element. Relative intensities still vary over the field of view with laser intensity, so a flatfield image was taken using free dye and used to correct for local intensity.

Point particles were picked from maximum intensity projections for each color channel using the Crocker and Grier algorithm and fit to 2D Gaussians to be filtered by size and intensity [19, 20]. Total intensity was integrated in a 2×2 pixel square region around particles sufficiently far from neighbors and local background was estimated from adjacent pixels and averaged over the duration of each video.

Slide specificity was measured by counting Cy5 surface localizations that photobleached in a single step per field of view (FOV) (Figure S4B). Competition experiments featured an additional 10X concentration of unlabeled competitor for surface binding sites (Figure S4C).

For FRET imaging, frames were alternated between 640nm and 532nm illumination to monitor the presence of both fluorophores. Traces were kept that showed simultaneous donor recovery upon acceptor bleaching and an observed γ factor of 1 ± 0.2. These were then filtered by intensity Cy5 emission intensity with 640nm excitation and Cy3 emission intensity upon 532 nm excitation after Cy5 photobleaching. Thresholds were set from the singly labeled protein-Cy3 and protein-Cy5 distributions to ensure single molecules of Cy3 and Cy5. The FRET ratio was calculated as the ratio of the acceptor emission to the total emission intensity for the trace prior to acceptor photobleaching. Traces with 10 or more time points exhibiting FRET were added to the overall distribution for a given condition. Empirical FRET was calculated as the ratio of Cy5 to total emission under 532 nm illumination.

For each FRET distribution, the Kernel Density Estimation (KDE) was calculated fit to the sum of two Gaussians with fitting parameters restricted to [0,∞) using the MATLAB curve fitting toolbox. Gaussian fits to Kernel Density Estimation of eFRET distributions were calculated using the MATLAB curve fitting toolbox to the function:

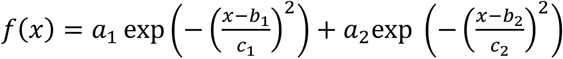

The relative area of the two Gaussians were calculated from the fitted values and the uncertainty was set by the minimum and maximum relative area within the 95% confidence interval of the fitting parameters.

Figures created with BioRender.com

